# Differences in external, internal oral and chondrocranial morphology of the tadpole of *Corythomantis greeningi* Boulenger, 1896 (Anura: Hylidae)

**DOI:** 10.1101/2020.10.02.324459

**Authors:** Lucas Rafael Uchôa, Claylton A. Costa, Antonia Joyce S. Santos, Rayone A. Silva, Felipe P. Sena, Etielle B. Andrade

## Abstract

The genus *Corythomantis* currently comprises a single species, *Corythomantis greeningi*, a hylid widely distributed in xerophilic and subhumid morphoclimatic regions of Brazil, mainly in the Northeast region. Recently the external morphology, internal oral anatomy, and chondrocranium of *C. greeningi* tadpoles were described from specimens collected in the state of Bahia, however, we observed some differences in morphology of individuals from the state of Piauí, northeastern Brazil. The tadpoles were collected during the 2019 rainy season and 14 individuals were used to describe and compare the larval characters. We observed differences in external, internal oral and chondrocranial morphology in relation to specimens previously described, especially in oral disc, number and shape of oral cavity papillae, and some chondrocranium structures, as: *cartilago suprarostralis*, *cornua trabeculae*, *fontanella frontoparietalis*, *cartilago orbitalis* e *planum hypobranchiale*. Our results point to the occurrence of heterochrony in *C. greeningi*, but we do not rule out the possibility that tadpoles belong to different species. Further studies involving a greater number of tadpoles at different stages, combined with genetic, acoustic, and morphological factors of adult specimens may establish the variation degree of *C. greeningi* in different regions of northeastern Brazil.

**RESUMO:** O gênero *Corythomantis* compreende atualmente uma única espécie, *Corythomantis greeningi*, um hilídeo amplamente distribuído nas regiões morfoclimáticas xerofílicas e subúmidas do Brasil, principalmente na região Nordeste. Recentimente foram descritas a morfologia externa, anatomia oral interna e condrocrânio do girino de *C. greeningi* a partir de espécimes coletados no estado da Bahia, no entanto, observamos algumas diferenças na morfologia dos indivíduos coletados na região norte do estado do Piauí, Nordeste do Brasil. Os girinos foram coletados durante o período chuvoso de 2019 e 14 indivíduos foram utilizados para descrição e comparação dos caracteres larvais. Observamos diferenças na morfologia externa, oral interna e no condrocranio do girino em relação ao descrito anteriormente, sobretudo no disco oral, no número e formato de papilas cavidade oral e algumas estruturas do condrocrânio, como: *cartilago suprarostralis*, *cornua trabeculae*, *fontanella frontoparietalis*, *cartilago orbitalis* e *planum hypobranchiale*. Nossos resultados apontam a ocorrência de heterocronia em *C. greeningi*, porém não descartamos a possibilidade dos girinos pertencerem a espécies diferentes. Estudos futuros envolvendo uma maior área de distribuição e maior número de indivíduos em estágios diferentes, aliados a fatores genéticos, acústico e morfológicos dos espécimes adultos poderão estabelecer o grau de variação de *C. greeningi* em diferentes regiões do Nordeste brasileiro.

## Introduction

Knowledge about the tadpole biology, especially related to morphology and anatomy, is an important source of information to understand taxonomic diversity, natural history, ecology of anuran species (Heyer *et al.*, 1990; Duellman and Trueb, 1994; Altig and Johnston, 1989; Altig and McDiarmid, 1999b). Since the amphibian larval morphology is directly related to evolutionary and ecological factors of the environment in which they live (Graham and Fine, 2008), their study guarantees a support for the understanding of natural patterns of species distribution, community structuring, morphological specialization, and even phylogenetic diversification (Lauder, 1981; Losos, 1990; Grosjean *et al.*, 2004).

Throughout the history of anurans phylogeny, larval characters have been used as a tool to clarify the systematics and evolution of the group (Lataste, 1879; Orton, 1953; Starrett, 1973; Sokol, 1975). Due to the high morphological diversity of tadpoles (Altig and McDiarmid, 1999a, b), some phylogenetic proposals for anurans were based exclusively on larval characters (Larson and de Sá, 1998; Haas, 2003) encompassing both broader taxonomic levels as species level (Larson and de Sá, 1998; Larson, 2005; d’Heursel and Haddad, 2007; Candioti, 2008).

The genus *Corythomantis* Boulenger, 1896, inserted in the subfamily Lophyohylinae (Frost, 2020), currently comprises a single species, *Corythomantis greeningi* Boulenger, 1896 (Blotto *et al.*, 2020). Belonging to the group of casque-head tree frogs, due to the total connection between skull bones and head mineralized dermis (Trueb, 1970; Jared *et al.*, 2005), *C. greeningi* is a hylid widely distributed in xerophilic and subhumid morphoclimatic regions of Brazil, mainly in the Northeast region (Frost, 2020).

Previous studies have described the external morphology of *C. greeningi* larvae, based on specimens collected in the municipalities of Feira de Santana and Morro do Chapéu (Juncá *et al.*, 2008), and internal oral anatomy and chondrocranium, from specimens collected in Barreira municipality, all in the state of Bahia (Oliveira *et al.*, 2017)(Fig. 1). However, we observed some differences in tadpoles collected in temporary streams in the northern region of the state of Piauí. Herein we describe differences found in external morphology, internal oral anatomy and chondrocranium of *C. greeningi* tadpoles from Pedro II municipality, state of Piauí, northeastern Brazil, and provide a brief commentary on larvae of the subfamily Lophyohylinae.

**Figure 1.**
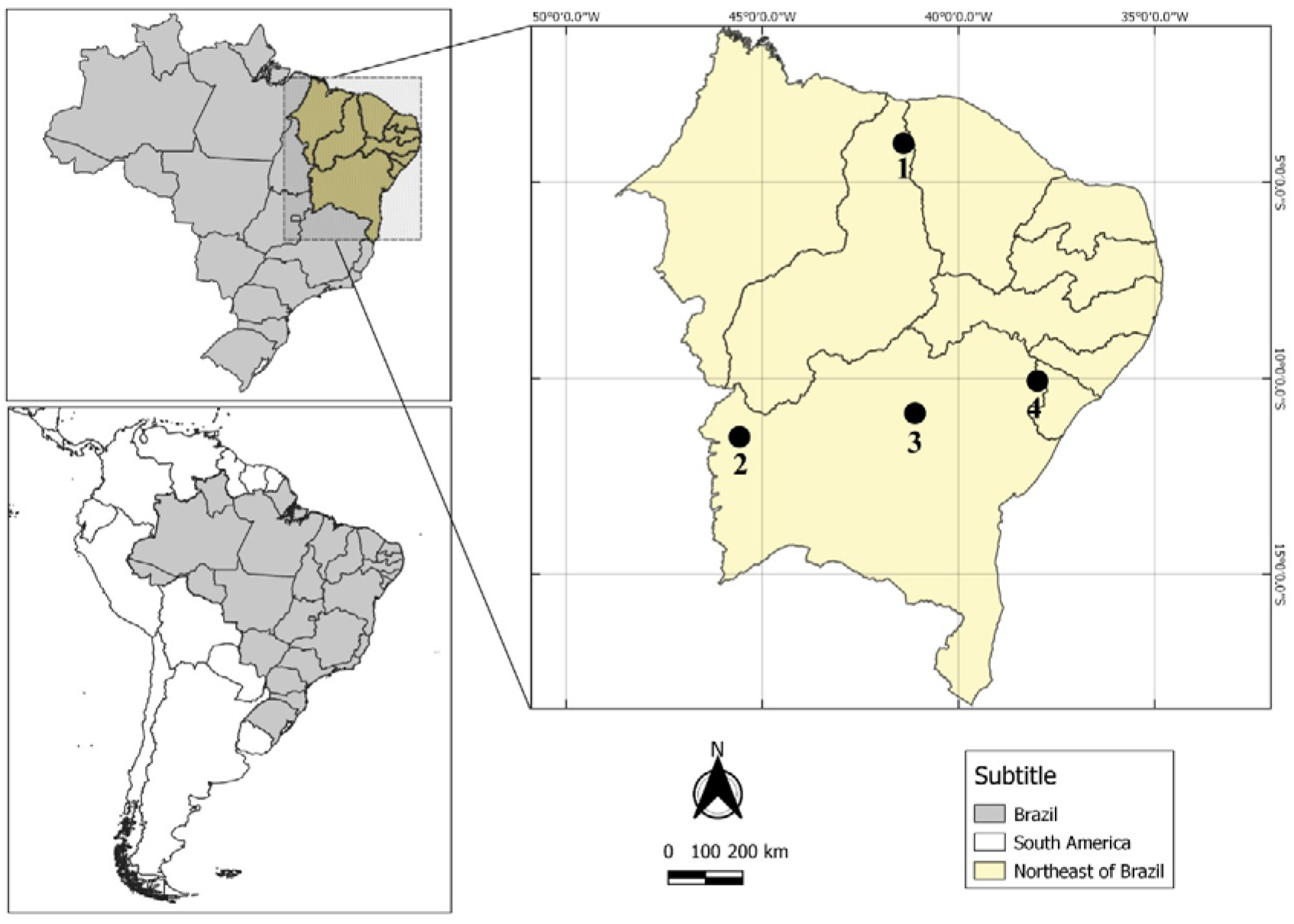
Collection points and literature records with description of *Corythomantis greening* tadpoles. 1 - Municipality of Pedro II, state of Piauí (present study); 2 - Barreiras (Oliveira *et al.*, 2017), 3 - Morro do Chapéu and 4 - Feira de Santana (Juncá *et al.*, 2008), state of Bahia.

## Materials and methods

Tadpoles were collected during the 2019 rainy season in temporary streams located in Pedro II municipality (4°30’34”S and 41°29’20”W, datum WGS84), northern region of the state of Piauí, northeastern Brazil (Fig. 1). The tadpoles were preserved in 4% formalin. Some specimens were raised in an aquarium until complete metamorphosis for correct identification. Vouchers specimens were deposited in the Biological Collection of the Instituto Federal de Educação, Ciência e Tecnologia do *Piauí* - IFPI *Campus* Pedro II (CBPII 99). Species identification was made based on morphological characters described by Juncá *et al.* (2008) and staged according to Gosner (1960).

External morphology description of *C. greeningi* tadpole was based on five stage 35 specimens (CBPII 100). External morphology followed Pezzuti (2011) and Andrade *et al.* (2018) and terminology followed Altig and McDiarmid (1999a) and Altig (2007). Sixteen morphometric measurements were taken: total length (TL), body length (BL), body width (BW), body height (BH), tail length (TaL), maximum tail height (MTH), tail musculature height (TMH), tail musculature width (TMW), dorsal fin height (DFH), ventral fin height (VFH), eye diameter (E), interorbital distance (IO), eye-nostril distance (END), internarial distance (IND), nostril-snout distance (NS) and oral disc width (ODW). Exclusively for the total length (TL), body length (BL), and tail length (TaL), we used a digital caliper with 0.01 mm accuracy. All other measures were taken using software TC Capture coupled to a stereoscopic microscope. All measurements (mean and standard deviation) are expressed in millimeters (Table 1).

**Table 1.**
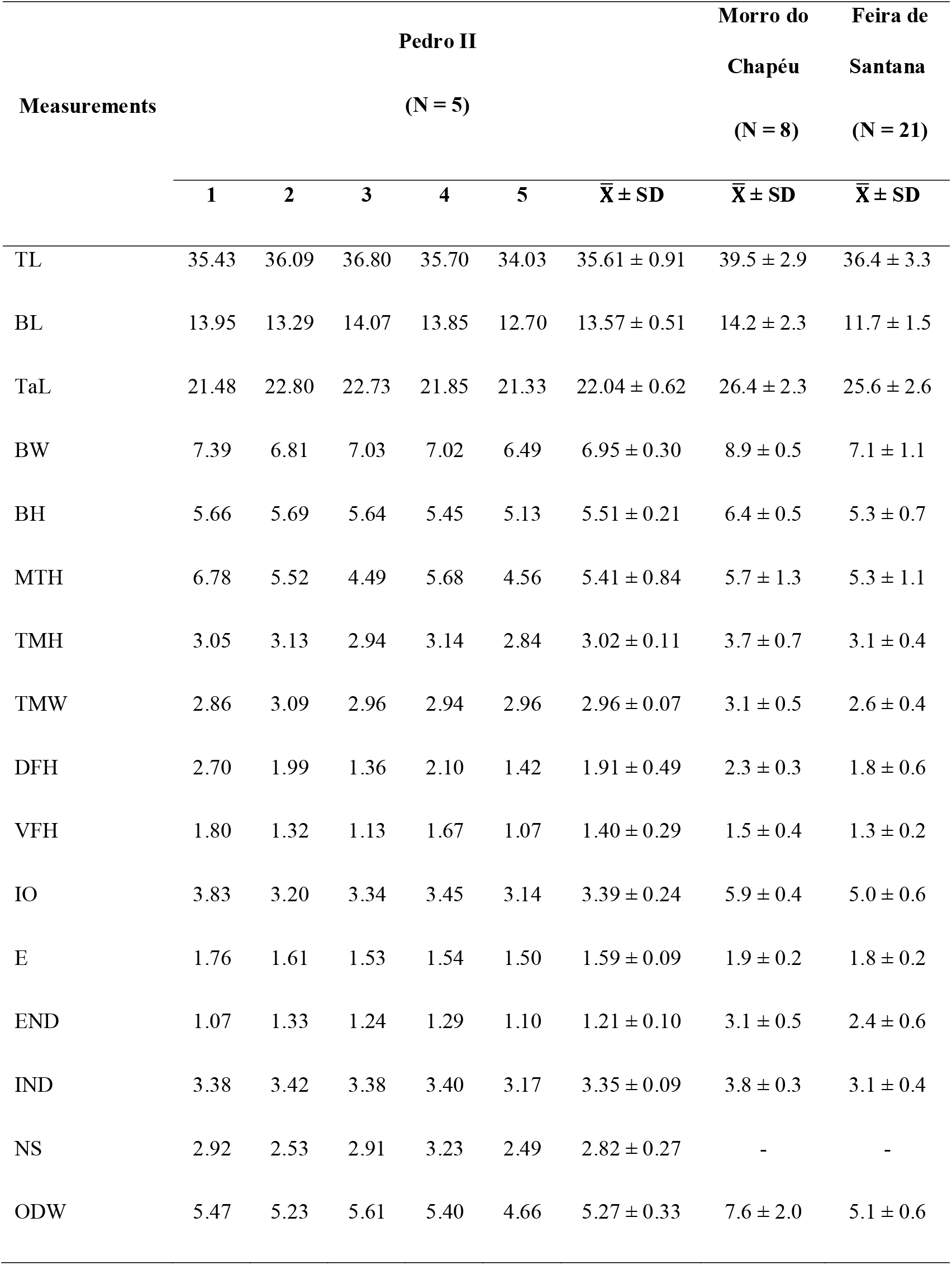
Measurements (in mm) of *Corythomantis greeningi* tadpoles (n = 5; stage 35) collected in Pedro II municipality, state of Piauí, and specimens (stage 34 and 36) collected in the municipalities of Feira de Santana and Morro do Chapéu, state of Bahia, northeastern Brazil (Juncá *et al.* 2008).

| Measurements | Pedro II<br>(N = 5) |  |  |  |  |  | Morro do<br>Chapéu<br>(N = 8) | Feira de<br>Santana<br>(N = 21) |
| --- | --- | --- | --- | --- | --- | --- | --- | --- |
| | 1 | 2 | 3 | 4 | 5 | $\bar{X} \pm SD$ | $\bar{X} \pm SD$ | $\bar{X} \pm SD$ |
| TL | 35.43 | 36.09 | 36.80 | 35.70 | 34.03 | 35.61 ± 0.91 | 39.5 ± 2.9 | 36.4 ± 3.3 |
| BL | 13.95 | 13.29 | 14.07 | 13.85 | 12.70 | 13.57 ± 0.51 | 14.2 ± 2.3 | 11.7 ± 1.5 |
| TaL | 21.48 | 22.80 | 22.73 | 21.85 | 21.33 | 22.04 ± 0.62 | 26.4 ± 2.3 | 25.6 ± 2.6 |
| BW | 7.39 | 6.81 | 7.03 | 7.02 | 6.49 | 6.95 ± 0.30 | 8.9 ± 0.5 | 7.1 ± 1.1 |
| BH | 5.66 | 5.69 | 5.64 | 5.45 | 5.13 | 5.51 ± 0.21 | 6.4 ± 0.5 | 5.3 ± 0.7 |
| MTH | 6.78 | 5.52 | 4.49 | 5.68 | 4.56 | 5.41 ± 0.84 | 5.7 ± 1.3 | 5.3 ± 1.1 |
| TMH | 3.05 | 3.13 | 2.94 | 3.14 | 2.84 | 3.02 ± 0.11 | 3.7 ± 0.7 | 3.1 ± 0.4 |
| TMW | 2.86 | 3.09 | 2.96 | 2.94 | 2.96 | 2.96 ± 0.07 | 3.1 ± 0.5 | 2.6 ± 0.4 |
| DFH | 2.70 | 1.99 | 1.36 | 2.10 | 1.42 | 1.91 ± 0.49 | 2.3 ± 0.3 | 1.8 ± 0.6 |
| VFH | 1.80 | 1.32 | 1.13 | 1.67 | 1.07 | 1.40 ± 0.29 | 1.5 ± 0.4 | 1.3 ± 0.2 |
| IO | 3.83 | 3.20 | 3.34 | 3.45 | 3.14 | 3.39 ± 0.24 | 5.9 ± 0.4 | 5.0 ± 0.6 |
| E | 1.76 | 1.61 | 1.53 | 1.54 | 1.50 | 1.59 ± 0.09 | 1.9 ± 0.2 | 1.8 ± 0.2 |
| END | 1.07 | 1.33 | 1.24 | 1.29 | 1.10 | 1.21 ± 0.10 | 3.1 ± 0.5 | 2.4 ± 0.6 |
| IND | 3.38 | 3.42 | 3.38 | 3.40 | 3.17 | 3.35 ± 0.09 | 3.8 ± 0.3 | 3.1 ± 0.4 |
| NS | 2.92 | 2.53 | 2.91 | 3.23 | 2.49 | 2.82 ± 0.27 | - | - |
| ODW | 5.47 | 5.23 | 5.61 | 5.40 | 4.66 | 5.27 ± 0.33 | 7.6 ± 2.0 | 5.1 ± 0.6 |

Five stage 36 tadpoles (CBPII 111) were prepared for analysis of internal oral anatomy according to Wassersug (1976). Internal oral anatomy terminology followed Wassersug (1976 and 1980) and Wassersug and Heyer (1988). Chondrocranium description was based on four tadpoles in stages 34 and 36 (CBPII 112), following Cannatella (1999), Haas (2003), Nascimento (2013), and Oliveira *et al.* (2017). The specimens were cleared and stained using the Taylor and Van Dyke (1985) technique with modifications. Chondrocranial terminology followed Larson and de Sá (1998), Cannatella (1999), and Oliveira *et al.* (2017).

## Results

### External Morphology

The tadpole of *C. greeningi* has an elliptical-elongated body (BW/BL = 51.21%) in dorsal view and depressed in lateral view (BH/BW = 79.28%), corresponding to approximately 38% of TL (Table 1; Fig. 2). Rounded snout in dorsal view and sloped in lateral view. Circular nostrils, located dorsally, with openings directed anterodorsolaterally, closer to eyes than snout (END/NS = 66.48%), without projections on the inner margins. Internarial distance approximately equal to interorbital distance. Dorsal eyes, dorsolaterally directed, representing about 12% of BL and 29% of BH. Interorbital distance about 49% of BH. Spiracle sinistral, cylindrical and short, positioned lateroventrally at the middle third of BL, with posterodorsal opening and visible in dorsal view. Spiracular opening free with the inner wall longer than the outer wall. Spiral intestinal tube with inflection point displaced from the abdomen center. Ventral tube medial, entirely fused to ventral fin, with slightly dextral opening. Medium height tail, corresponding about 62% of TL, and presenting an acute termination. Tail musculature robust, presenting a height about 55% of BH and width about 42% of BW, with sharp tapering from the anterior third of the tail. Dorsal fin of medium height, with margin slightly convex, arising near the body-tail junction. Dorsal fin higher than the ventral fin (VFH/DFH = 73.30%), with maximum height located in the middle third of the tail. Ventral fin of medium height, originating at the level of the ventral tube.

**Figure 2.**
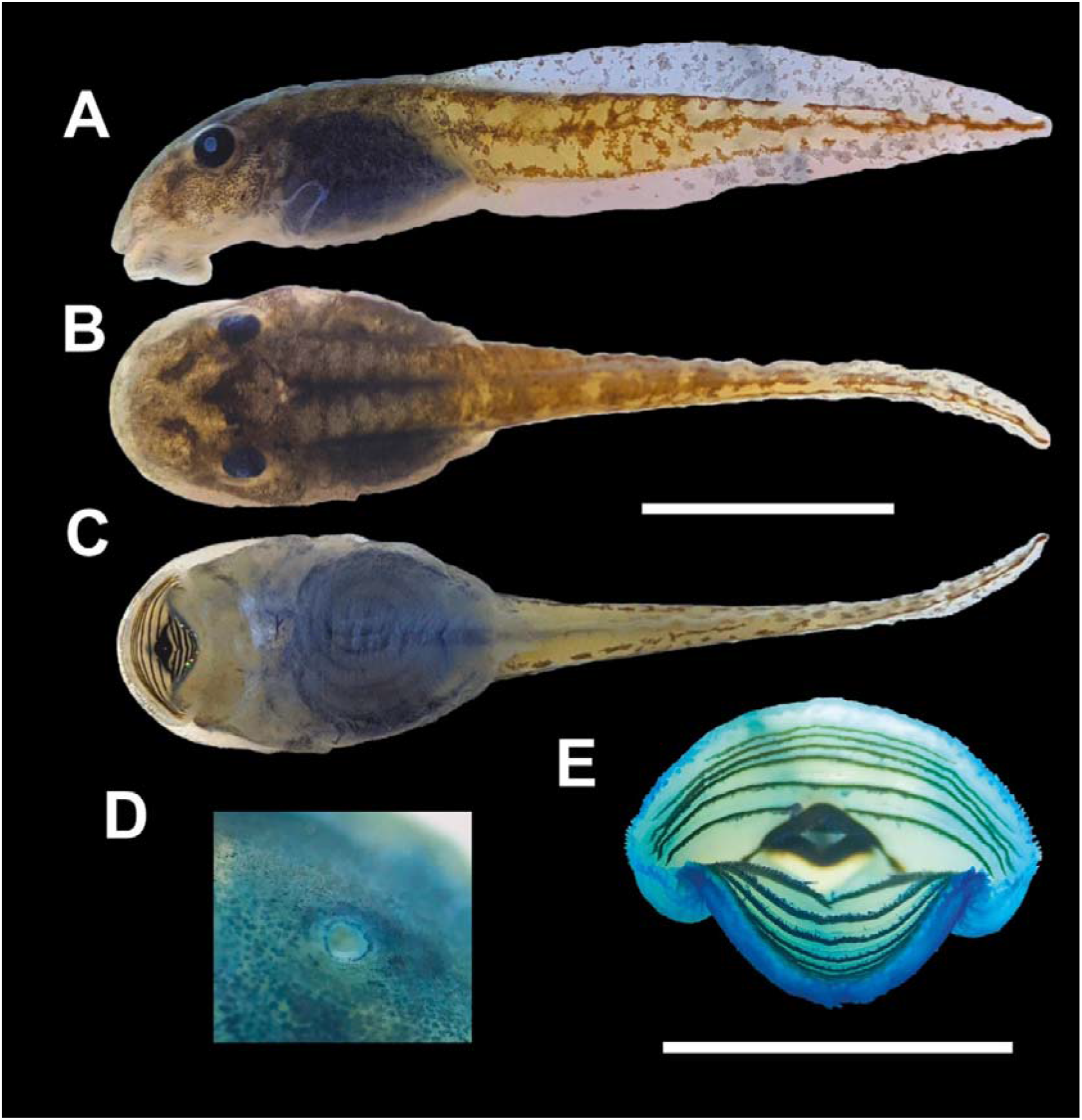
*Corythomantis greeningi* tadpole collected in Pedro II municipality, state of Piauí, northeastern Brazil. Specimens at stage 35 (CBPII 100). (A) lateral, (B) dorsal and (C) ventral views, (D) nostril and (E) oral disc. Scale bar = 5 mm.

Oral disk is common with the presence of keratinized structures (Fig. 2E), emarginated and positioned ventrally. It presents a row of marginal papillae uniseriate without interruptions. Submarginal papillae are present on the posterior lip and in smaller numbers on the sides of the oral disc, and absent on the anterior lip. Labial tooth row formula LTRF 6(6)/8(1), with A1<A2<A3=A4=A5=A6 and P1=P2=P3=P4=P5=P6>P7>P8. Denticles absent on the sides of the oral disc. Lower jaw V-shaped and the upper jaw triangular, both serrated with a wide base.

### Internal oral morphology

Buccal floor (Fig. 3A) diamond-shaped, slightly wider than long (length/width = 87.5%). Two pairs of overlapping infralabial papillae are present, the upper pair is larger and hand-shaped, and the lower pair digitiform. Lingual bud well developed, with a pair of long and tapering lingual papillae. Each lingual papillae has small lateral projections along its structure. Buccal pockets are large, deep, and transversely oriented towards the buccal floor arena, with the presence of 11–12 prepocket papillae digitiform varying in size on each side, being 4–5 papillae fused at the base. Buccal floor arena (bfa) with 22–26 digitiform papillae varying in size on each side, the largest being located laterally in the floor arena and the smallest in the central region. Some pustules are present, located mainly in the posterior region of the floor arena near the glottis. Wide ventral velum with three marginal projections on each side separated by a well-marked median notch. Well-defined secretory region, with distinct and exposed glottis.

**Figure 3.**
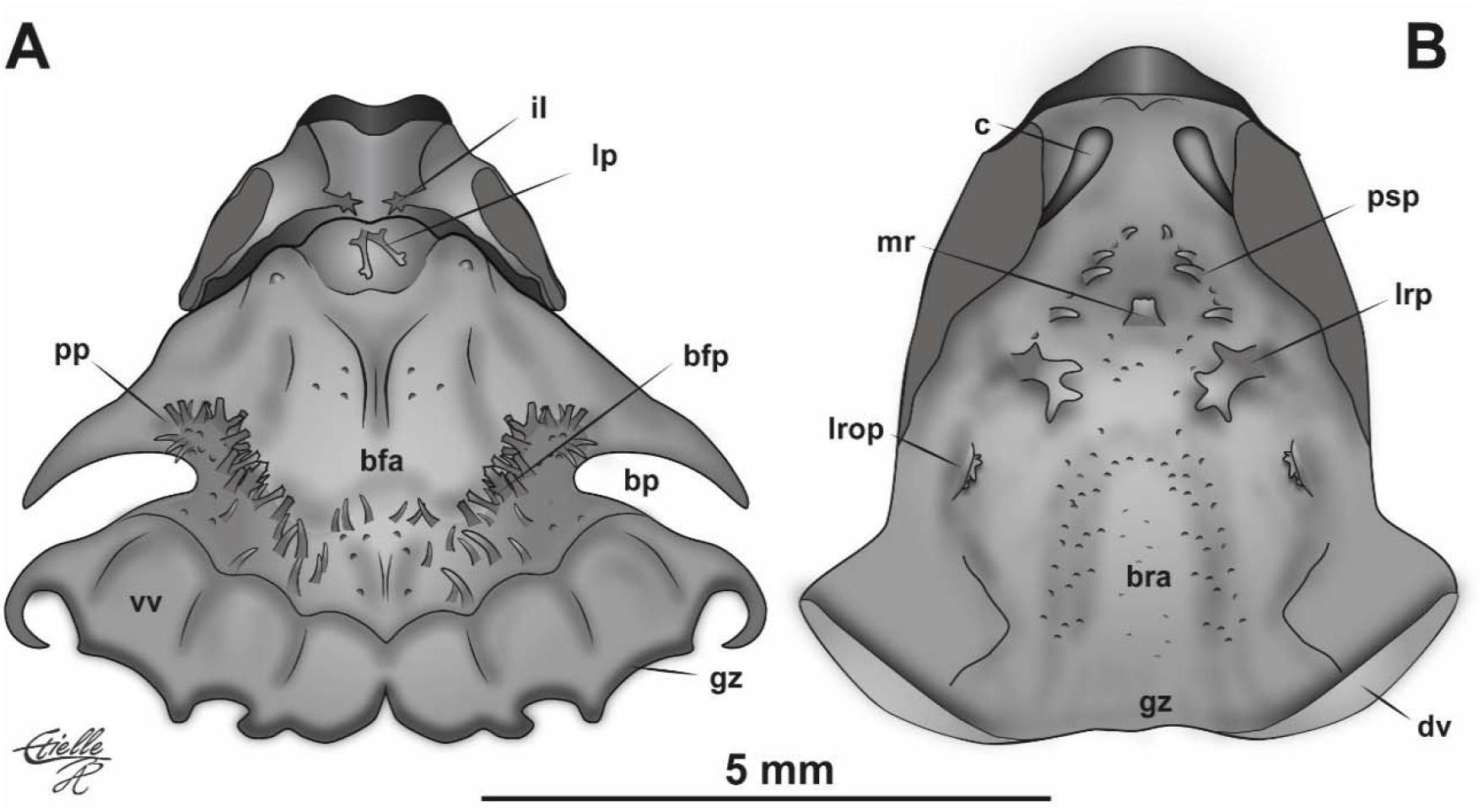
Internal oral anatomy of *Corythomantis greeningi* (Stage 36; CBPII 111) tadpole. **(A)** Buccal floor and **(B)** buccal roof. Abbreviations: bfa: buccal floor arena; bfp: buccal floor arena papillae; bp: buccal pocket; bra: buccal roof arena; c: choanae; dv: dorsal velum; gz: grandular zone; il: infralabial papillae; lp: lingual papillae; lrop: lateral roof papillae; lrp: lateral ridge papillae; mr: median ridge; pp: prepocket papillae; psp: postnarial papillae; vv: ventral velum. Scale bar = 5 mm.

Buccal roof (Fig. 3B) is overall triangular. Prenarial arena is wide and concave, containing a Y-shaped ridge with irregular margins. Narrow choanae, medially curved and longitudinally oriented towards the prenarial arena. Postnarial arena has two rows of 2–6 conical papillae on each side, arranged in the inverted V-shaped. Median ridge is small and overall trapezoidal, with a narrow base and serrated upper margin. The lateral ridge papillae are well developed, broad-based, hand-shaped with four to five projections on the anterior border, and with the presence of 2–3 small conical papillae close to their base. The buccal roof arena (bra) without papillae and with the presence of some pustules evenly distributed. About 4–5 lateral papillae are found aligned on each side of the buccal roof. The lateral papillae are conical, with a rounded apex, and oriented towards the midline. Glandular zone is distinct. Dorsal velum is wide laterally, with a folded glandular border.

### Chondrocranial Morphology

The chondrocranium is elliptical, slightly longer than wide (width/length = 86%), and depressed in lateral view (height/width = 53%), being wider at the level of *arcus subocularis* and higher at the level of *cornua trabeculae* (Fig. 4).

**Figure 4.**
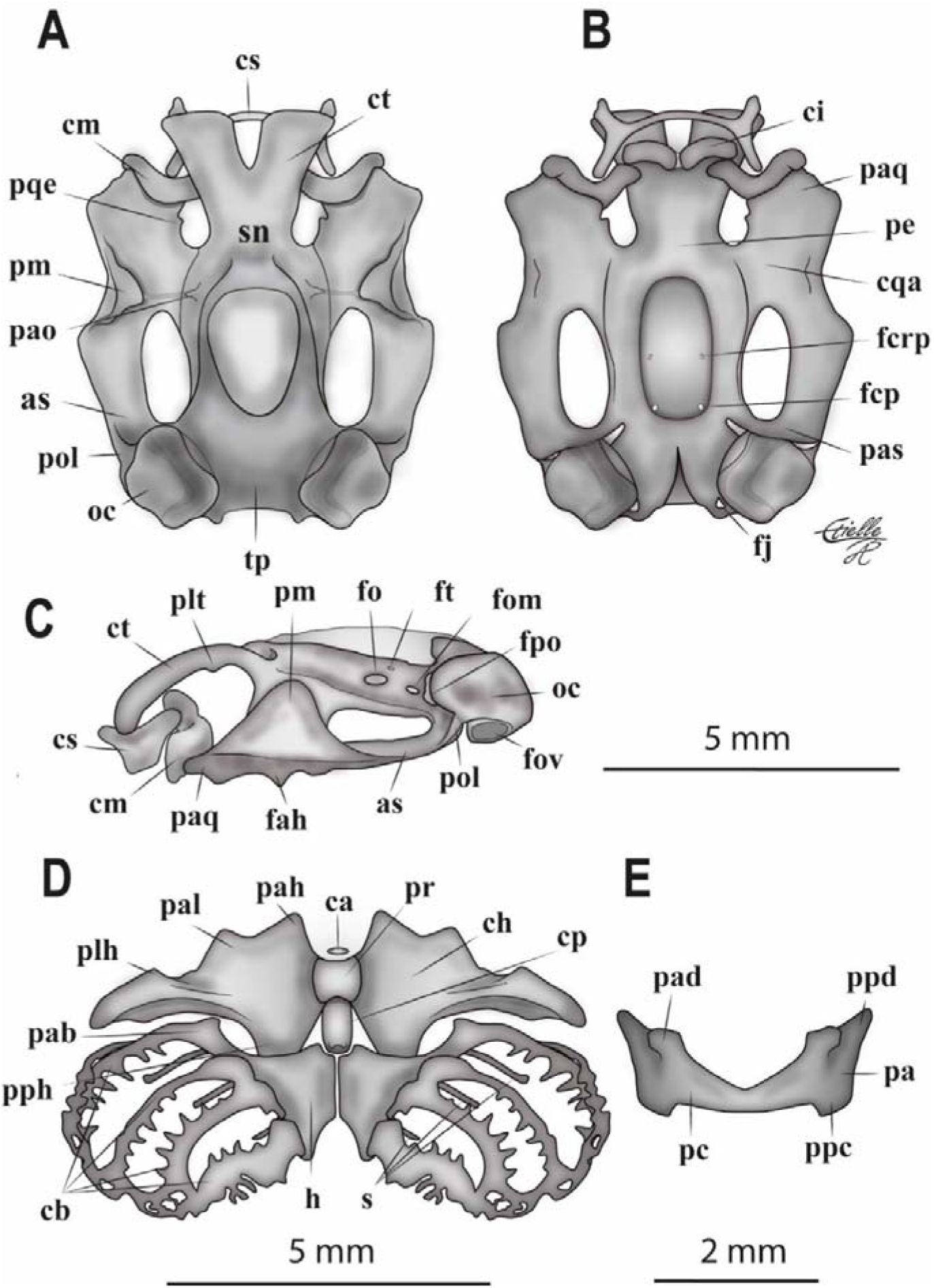
Chondrocranium of *Corythomantis greeningi* tadpole collected in the Pedro II municipality, state of Piauí, northeastern Brazil, at stage 36 (CBPII 112). (A) dorsal, (B) ventral, (C) lateral views, (D) ventral view of the hyobranchial apparatus, and (E) frontal view of the *cartilago suprarostralis*. Abbreviations: as - *arcus subocularis*; ca - *copula anterior*; cb - *ceratobranchiales*; ch - *ceratobranchiales*; ci - *cartilago infrarostralis*; cm - *cartilago Meckeli*; cqa - *comissura quadratocranialis anterior*; cp - *copula posterior*; cs - *cartilago suprarostralis*; ct - *cornua trabeculae*; fah - *facies articularis hyalis*; fcp - *foramen caroticum primarium*; fcrp - *foramen craniopalatinum*; fj - *foramen jugulare*; fo - *foramen opticum*; fom - *foramen oculomotorium*; fov - *fenestra ovalis*; fpo - *foramen prooticum*; ft - *foramen trochleare*; h - *hypobranchiale*; oc - otic capsule; pa - *pars alaris*; pab - *processus anterior branchialis*; pad - *processus anterior dorsalis*; pah - *processus anterior hyalis*; pal - *processus anterolateralis hyalis*; pao - *processus antorbitalis*; paq - *processus articularis*; pas - *processus ascendens*; pc - *pars corporis*; pe - *planum ethmoidale*; plh - *processus lateralis hyalis*; plt - *processus lateralis trabeculae*; pm - *processus muscularis quadrati*; pol - *processus oticus larval*; ppc - *processus posterior ventralis*; ppd - *processus posterior dorsalis*; pph - *processus posterior hyalis*; pqe - *processus quadratoethmoidalis*; s - *spicula*; sn - *septum nasi*; and tp - *tectum parientalis.*

#### Ethmoidal Region

The *cartilago suprarostralis* consists of *pars corporis* and *pars alaris*. *Par corporis* of *cartilago suprarostralis* is rectangular, ventrally fused, with a wide V-shaped notch. *Par alaris* wide and rectangular, fully fused to the *par corporis*, with a flat ventral surface. *Par alaris* has a long and acute *processus anterior dorsalis*, exceeding the anterior margin of the *cornua trabeculae*, and a long, rounded *processus posterior dorsalis*. *Cornua trabeculae* are short, robust, and ventrally curved, distally divergent in a V-shaped, presenting a V-shaped notch in its distal margin. *Processus lateralis trabeculae* are present, short, and located close to the *planum ethmoidale*. *Planum ethmoidale* broad dorsally, anteriorly delimited by *taenia tecti ethmoidales*, dorsolaterally by *taenia tecti marginalis* and posteriorly by *tectum parientalis*, defining a reduced and undivided *fenestra frontoparietalis*. *Lamina orbitonasalis* well developed with the presence of *foramen orbitonasalis*.

#### Orbitotemporal Region

*Planum intertrabeculare* appears as a thin and slightly chondrified leaf, which closes the *fenestra basicranialis*, forming the central area of the cranial floor. *Foramen caroticum primarium* and *foramen craniopalatinum* are present, the first being larger than the second. *Cartilago orbitalis* well chondrified, allowing the visualization of four foramina: *foramen opticum* broad and elliptical, *foramen trochleare* smaller and narrow located just above the *foramen opticum*, *foramen opticum* medium and elliptical, medially located between *foramen prooticum* and *foramen opticum*, and *foramen prooticum* located between the anterior margin of the optic capsules and *pila antotica*.

#### Palatoquadrate

Long cartilage with smooth margins connected to the braincase through the *processus articularis quadrati*, *processus ascendens*, and *processus oticus larval*. *Processus ascendens* short, broad, and attached to the *pila antotica*. *Arcus subocularis* robust, posteriorly narrower. *Fenestra subocularis* ovoid, longer than broad. *Processus articularis quadrati* short, wider than long, articulating anteriorly with *cartilago Meckeli*. *Processus muscularis quadrati* broad and triangular, curves medially towards the braincase and joins a small *processus antorbitalis* of the *planum ethmoidale* through the barely visible *ligamentum tectum*. A small and triangular *processus quadratoethmoidalis* is present, located on the inner margin of the broad *commissura quadratocranialis anterior*. *Processus pseudopterygoideus* absent. *Facies articularis hyalis* triangular, located below the *processus muscularis quadrati*, articulating with ceratohyal.

#### Otoocipital Region

Otic capsules are rhomboid corresponding to about 20% of total chondrocranial length. On the lateral wall of the otic capsules, a *processus anterolateralis* of the larval parotic crest protrudes horizontally and connects to the posterior curvature of palatoquadrate forming a *processus oticus larval*. Otic capsules are connected dorsally to each other by *tectum parientalis*, forming the dorsal roof of *foramen magnum*. *Arcus occipitalis* extends posteromedially to the otic capsules from the *planum basale*, forming the medial and ventral margins of the *foramen jugulare* and occipital condyles. A small *foramen perilymphaticum inferior* can be noticed in the ventromedial surface of the optic capsule. *Fenestra ovalis* of moderate size located ventrolaterally just below the larval parotic crest.

#### Cartilago Meckeli

Lower jaw is formed by *cartilago Meckeli* and *cartilago infrarrostralis*, representing about 72% of the width of the chondrocranium. *Cartilago Meckeli* are sigmoid located ventrally to the *cornua trabeculae* and articulates dorsolaterally with the *processus articularis quadrati* through the short *processus retroarticularis*. *Processus dorsomedialis* and *processus ventromedialis* of *cartilago Meckeli* support the *cartilago infrarostralis*, in which they are medially connected by the *commissura intermandibularis*. *Cartilago infrarostralis* are slightly curved, located medioventrally to the *cartilago Meckeli* and ventrally to the *cornua trabeculae*, and positioned almost perpendicular to the main axis of the chondrocranium.

#### Hyobranchial Apparatus

*Ceratohyalia* are broad and subtriangular, medially flat, oriented perpendicular to the main axis of the chondrocranium. In lateral and dorsal view there is a vertical condylar expansion, the *processus articularis*, which articulates with the *facies articularis hyalis* of the *palatoquadrate*. Anteriorly, each margin of *ceratohyalia* has a well-developed *processus anterior hyalis*, triangular and laterally curved, a *processus anterolateralis hyalis*, triangular, smaller and slightly wider, and *processus lateralis hyalis*, more discreet than the others. In some individuals, it is possible to observe a small elevation between the *processus anterolateralis hyalis* and *processus lateralis hyalis*. *Condylus articularis* is long. Posteriorly, *ceratohyalia* has a well-developed *processus posterior hyalis*.

*Ceratohyalia* are medially connected to a rounded and slightly chondrified *pars reuniens*. *Copula anterior* is a small and elliptical cartilage transversely positioned over the *pars reuniens. Copula posterior* is rectangular and robust, presenting a short *processus urobrancialis*. *Copula posterior* connects posteriorly to the *planum hypobranchiale*, which are well-developed, broad, and triangular cartilaginous plaques that support the branchial baskets. *Planum hypobranchial*e are medially articulated by the *commissura inter-hyal* diverging posteriorly in an inverted U-shaped with rounded edges. *Ceratobranchial I* continuous with the *planum hypobranchiale*. *Processus anterior branchialis* well-developed. *Ceratobranchiales* are joined distally by *commissura terminalis*. *Ceratobranchiales II*, *III*, and *IV* are syndesmotically connected to the *planum hypobranchiale*. Spicule I, II, and III are well-developed.

## Discussion

The subfamily Lophyohylinae has a great diversity in larval morphology of its species and different biological traits associated with oophagy and anti-predatory mechanisms (Blotto *et al.*, 2020). Of the 88 species of Lophyohylinae currently recognized approximately 44.3% (39 species) have information about their larvae: *C. greeningi*, *Itapotihyla langsdorffii* (Duméril and Bibron, 1841) (Pimenta and Canedo, 2007), *Nyctimantis arapapa* Pimenta, Napoli and Haddad, 2009 (Lourenço-de-Moraes *et al.*, 2013), *N. brunoi* Miranda-Ribeiro, 1920 (Wogel *et al.*, 2006), *N. siemersi* (Mertens, 1937) (Cajade *et al.*, 2010), *Osteocephalus buckleyi* (Boulenger, 1882) (Hero, 1990), *O. cabrerai* (Cochran and Goin, 1970) (Menin *et al.*, 2011), *O. festae* (Peracca, 1904) (Ron *et al.*, 2010), *O. mimeticus* (Melin, 1941) (Henle, 1981), *O. oophagus* Jungfer and Schiesari, 1995, *O. taurinus* Steindachner, 1862 (Schiesari *et al.*, 1996), *O. verruciger* (Werner, 1901) (Ron *et al.*, 2010), *Osteopilus crucialis* (Harlan, 1826), *Os. dominicensis* (Tschudi, 1838), *Os. marianae* (Dunn, 1926) (Galvis *et al.*, 2014), *Os. ocellatus* (Linnaeus, 1758) (Lannoo *et al.*, 1987), *Os. pulchrilineatus* (Cope, 1870), *Os. septentrionalis* (Duméril and Bibron, 1841), *Os. vastus* (Cope, 1871), *Os. wilderi* (Dunn, 1925) (Galvis *et al.*, 2014), *Phyllodytes acuminatus* Bokermann, 1966 (Campos *et al.*, 2014), *P. brevirostris* Peixoto and Cruz, 1988 (Vieira *et al.*, 2009), *P. edelmoi* Peixoto, Caramaschi and Freire, 2003, *P. gyrinaethes* Peixoto, Caramaschi and Freire, 2003 (Peixoto *et al.*, 2003), *P. luteolus* (Wied-Neuwied, 1821) (Campos *et al.*, 2014), *P. magnus* Dias, Novaes-e-Fagundes, Mollo, Zina, Garcia, Recoder, Vechio, Rodrigues and Solé, 2020 (Dias *et al.*, 2020), *P. melanomystax* Caramaschi, Silva and Britto-Pereira, 1992 (Caramaschi *et al.*, 1992), *P. praeceptor* Orrico, Dias and Marciano, 2018 (Santos *et al.*, 2019), *P. tuberculosus* Bokermann, 1966 (Campos *et al.*, 2014), *P. wuchereri* (Peters, 1873) (Magalhães *et al.*, 2015), *Tepuihyla obscura* Kok, Ratz, Tegelaar, Aubret and Means, 2015 (Kok *et al.*, 2015), *Trachycephalus atlas* Bokermann, 1966 (Barreto *et al.*, 2015), *T. coriaceus* (Peters, 1867) (Schiesari *et al.*, 1996), *T. cunauaru* Gordo, Toledo, Suárez, Kawashita-Ribeiro, Ávila, Morais and Nunes, 2013 (Grillitsch, 1992), *T. jordani* (Stejneger and Test, 1891) (Mcdiarmid and Altig 1990), *T. mesophaeus* (Hensel, 1867) (Prado *et al.*, 2003), *T. nigromaculatus* Tschudi, 1838 (Wogel *et al.*, 2000), *T. resinifictrix* (Goeldi, 1907) (Schiesari *et al.*, 1996), and *T. typhonius* (Linnaeus, 1758) (Schiesari *et al.*, 1996). Information about the external morphology of tadpoles is presented for all the species described above, however, few of them have the internal oral anatomy and chondrocranium described. Only *N. brunoi*, *C. greeningi*, and *T. typhonius* (approximately 3.5%) have the three detailed morphological descriptions for the tadpole, which limits the comparisons between the species belonging to the subfamily.

Recent molecular analysis between populations of *C. greeningi* from the states of Alagoas and Tocantins showed polyphyletism within the genus, indicating the need for further taxonomic studies involving the monotypic genus (Blotto *et al.*, 2020). Juncá *et al.* (2008) described the *C. greeningi* tadpole based on specimens from two different locations in the state of Bahia, reporting the occurrence of a dwarf population in Feira de Santana municipality. We observed a slightly smaller average body size (35.61 ± 0.91) for *C. greeningi* larvae registered in the northern region of Piauí when compared to populations registered in the municipality of Feira de Santana (36.4 ± 3.3) and Morro do Chapéu (39.5 ± 2.9), both in the state of Bahia (Juncá *et al.*, 2008). Although the specimens registered here are within the body variation range of the tadpoles from Bahia, it is not possible to make an accurate comparison since in the original description were used tadpoles in different stages (34-36), resulting in a greater amplitude in body size. Juncá *et al.* (2008) suggest that the smaller body size of the tadpoles from the municipality of Feira de Santana - BA is caused by anthropic factors, as such change in shelters quality for tadpoles and food availability, or by acceleration of metamorphosis in water bodies with short hydroperiod. Phenotypic differences related to the tadpoles morphological characters are well documented in the literature, including among populations of the same species (Zhao *et al.*, 2017; Jordani *et al.*, 2019), since anuran larvae are highly sensitive both to the physical environment as to their biotic interactions regarding trophic specializations (Eterovick *et al.*, 2010; Michel, 2012; Zhao *et al.*, 2014, Johnson *et al.*, 2015).

Some small differences were observed (tadpole characteristics from Bahia in parentheses): body elliptical-elongated in dorsal view (oval body), circular nostrils (oval), body about 38% of TL (36% of TL), IO = IND (IO > IND), tail muscle tapered in the posterior third (slightly tapered), TMW about 42% of BW (35% of BW), intestinal tube inflection point displaced from the abdomen center (inflection point located in the abdomen center). Besides, we observed in all the tadpoles a labial tooth row formula (LTRF) = 6(6)/8(1) and a lot of papillae submarginal on the lower labium, differing from those recorded in the tadpole from Bahia (Juncá *et al.*, 2008). Oral disc morphological characteristics of *C. greeningi* are related to lotic watercourses, in which are adapted morphologically for suction (McDiarmid and Altig, 1999; Juncá *et al.*, 2008). The other structures were similar to those described by Juncá *et al.* (2008), with only minor morphometric variations. Recently, Dubeux *et al.* (2020) presented information about the external morphology of *C. greeningi* tadpoles from states of Alagoas and Rio Grande do Norte, which were similar to those presented here.

Regarding the internal oral anatomy of Lophyohylinae tadpole, only 15% of the species are described, and for some of them, only illustrations without detailed description are provided: *C. greeningi* (Oliveira *et al.*, 2017), *N. brunoi* (Wogel *et al.*, 2006), *N. siemersi* (Cajade *et al.*, 2010), *O. oophagus*, *O. taurinus*, *Os. septentrionalis* (Schiesari *et al.*, 1996), *Os. ocellatus* (Lannoo *et al.*, 1987), *P. brevirostris* (Vieira *et al.*, 2009), *P. wuchereri* (Magalhães *et al.*, 2015), *T. atlas* (Barreto *et al.*, 2015), *T. cunauaru* (Grillitsch, 1992), *T. resinifictrix* (Schiesari *et al.*, 1996), and *T. typhonius* (Schiesari *et al.*, 1996). We observed significative differences in the internal oral anatomy between the tadpoles from the states of Piauí and Bahia, mainly in the shape and number of papillae on the floor and buccal roof. Oliveira *et al.* (2017) did not provide details on the infralabial and lingual papillae shape, but observing the images of the oral cavity presented by authors, it is possible to notice differences in the shape of lingual papillae between specimens from Piauí (long and with projections) and Bahia (small, conical and without projections). In addition, the shape of lingual papillae found in *C. greeningi* tadpoles presented here does not resemble any other tadpole in the subfamily, since when present, the lingual papillae are simple and without lateral projections. The infralabial papillae act as respiratory, sensory, or mechanical structures (Wassersug, 1980), varying in number within the subfamily: absent in *Os. ocellatus* (Lannoo *et al.*, 1987); a pair in *N. brunoi* (Wogel *et al.*, 2006), *N. siemersi* (Cajade *et al.*, 2010), *O. oophagus*, *O. taurinus* (Schiesari *et al.*, 1996), *P. brevirostris* (Vieira *et al.*, 2009), and *P. wuchereri* (Magalhães *et al.*, 2015); and two pairs in *C. greeningi* (Oliveira *et al.*, 2017, present work), *Os. septentrionalis* (Lannoo *et al.*, 1987), *T. atlas* (Barreto *et al.*, 2015), *T. cunauaru* (Grillitsch, 1992), *T. resinifictrix* (Schiesari *et al.*, 1996), and *T. typhonius* (Schiesari *et al.*, 1996).

The high number of papillae observed on the buccal floor arena also differs from all species of the subfamily, since the maximum number of papillae recorded so far (13–15 papillae on each side) was found in *O. taurinus* and *O. oophagus* (Schiesari *et al.*, 1996). The specimens from Piauí have greater complexity concerning internal oral characters, and the large number of papillae in the buccal floor arena is consistent with species adapted to lotic environments (Wassersug, 1980), diverging from the results by Oliveira *et al.* (2017). These authors affirm that the *C. greeningi* tadpoles, although they have been found in lotic environments, are mainly similar to species adapted to lentic environments (Oliveira *et al.*, 2017). The buccal roof also showed marked differences, especially in the choanae direction, number of papillae in post-choanal arena, and median ridge shape. According to these characteristics, the population of *C. greeningi* in northern Piauí is mainly similar to *O. taurinus* and *T. cunauaru* (Grillitsch, 1992; Schiesari *et al.*, 1996). Typically, Lophyohylinae has a semicircular median crest (Schiesari *et al.*, 1996; Cajade *et al.*, 2010; Barreto *et al.*, 2015; Magalhães *et al.*, 2015; Oliveira *et al.*, 2017), but *C. greeningi* (populations from Piauí) diverges of this pattern by presents a trapezoid median crest, similar to *O. oophagus*, *T. cunauaru*, *T. resinifictrix* (Grillitsch, 1992; Schiesari *et al.*, 1996). Oliveira *et al.* (2017) report variation in median ridge (semicircular and trapezoidal), however, we observed no variation in the analyzed tadpoles.

Except for *O. ocellatus*, *O. septentrionalis*, *P. brevirostris*, and *P. wuchereri* (Lannoo *et al.*, 1987; Vieira *et al.*, 2009; Magalhães *et al.*, 2015), the typical shape of lateral ridge papillae is triangular with an irregular anterior margin (Grillitsch, 1992; Schiesari *et al.*, 1996; Cajade *et al.*, 2010; Barreto *et al.*, 2015; Oliveira *et al.*, 2017, present work). We observed a differentiated pattern, in which there are well-developed projections on the anterior margin of the lateral ridge papillae. Lateral roof papillae are common among the subfamily species, except in *O. septentrionalis* (Lannoo *et al.*, 1987), despite the variable number (Lannoo *et al.*, 1987, Schiesari *et al.*, 1996; Cajade *et al.*, 2010, Wogel *et al.*, 2006; Oliveira *et al.*, 2017; present work).

Only six species of Lophyohylinae have some type of information about the chondrocranium, which represents about 7% of the species. Among these species, *C. greeningi* (Oliveira *et al.*, 2017), *N. brunoi* (da Silva, 1994), and *P. gyrinaethes* (Candioti *et al.*, 2016) have a detailed description of the chondrocranium, while in *Os. ocellatus* (Lannoo *et al.*, 1987), *T. typhonius* (Fabrezi and Vera, 1997), *T. resinifictrix* (Haas, 2003) only a few structures are mentioned. Due to a lack of information on the Lophyohylinae chondrocranium, systematic comparisons of the structures become difficult. In addition, since these are species with different life histories subject to different ecological pressures (ecomorphology), there is a great variation among the chondrocranium already described (Oliveira *et al.*, 2017).

About chondrocranium, we observe differences among specimens from Piauí and those from Bahia (characters inside the parentheses): chondrocranium global shape (oval), the *cartilago suprarostralis* shape (*processus anterior dorsalis* short), notch shape between the *cornua trabeculae* (U shape), presence of the *processus lateralis trabeculae* (absent), presence of the *processus quadratoethmoidalis* (absent), reduced *fontanella frontoparietalis* (*fontanella frontoparietalis* large), presence of four foramina in the *cartilago orbitalis* wall (not visible), presence of a small *processus antorbitalis* (absent). Regarding the hyobranchial apparatus, the specimens differ overall by: *pars reuniens* shape (semicircular), *condylus articularis* size (short), *planum hypobranchiale* shape (narrow), and its posterior notch (inverted V-shaped). Chondrocranial morphology is very conserved and phylogenetically informative in phylogenetic hypotheses construction, even among closely related species (Larson and de Sá, 1998; Haas, 2003; Fabrezi and Quinzio, 2008). However, heterochronic variation in appearance and larval traits development can occur in some species (Larson, 2002; Fabrezi and Goldberg, 2009), which may explain the differences found in the *C. greeningi* chondrocranium. Nevertheless, based on *C. greeningi* polyphyly (Blotto *et al.*, 2020), we do not rule out the possibility that specimens from Piauí (present study) and those from Bahia (Juncá *et al.*, 2008; Oliveira *et al.*, 2017) belong to different species (Marques *et al.*, 2019).

In general, the chondrocranium of *C. greeningi* is quite similar to *N. brunoi* and *T. typhonius* by presents ovoid or elliptical shape, a rectangular and ventrally fused *cartilago suprarostralis*, wide *cornua trabeculae*, robust and well-developed *processus muscularis quadrati*, and presence of *processus oticus larval* (da Silva, 1994; Fabrezi and Vera, 1997; Oliveira *et al.*, 2017; present work) and differs completely from *P. gyrinaethes* (Candioti *et al.*, 2016), corroborating the phylogenetic tree of Blotto *et al.* (2020). According to Oliveira *et al.* (2017), the tadpoles of *C. greeningi* are more similar to Pelodryadinae tadpoles, specialized in shaving the bottom of lotic environments. However, the Hylidae and Pelodryadidae families diverged in the Paleocene (about 61.8 Ma; Duellman *et al.*, 2016) indicating that the tadpoles specialized in suction evolved several times independently, guided mainly by ecological aspects of the natural environments (Haas and Richards, 1998).

Our results show marked differences in external morphology, internal oral anatomy, and chondrocranium between *C. greeningi* tadpoles from the states of Bahia and Piauí, especially in the oral disc, number and papillae shape in the oral cavity, and some chondrocranium structures. Future studies involving a larger number of individuals at different stages and collected across the species range will be essential to establish these differences as population variations. Besides, broader studies on genetic, acoustic, and morphological factors of adult specimens may establish the degree of variation of *C. greeningi* in different regions of Northeast Brazil.

## Acknowledgements

We thank the Instituto Federal de Educação, Ciência e Tecnologia do Piauí - IFPI for providing a grant through the Programa de Apoio à Pesquisa, Estruturação e Reestruturação Laboratorial - PROAGRUPAR-INFRA (edital n° 077 de 07/05/2018), and to Instituto Chico Mendes de Conservação à Biodiversidade by colletion licence (#61838-2/19).

## Notes

### Competing Interest Statement

The authors have declared no competing interest.

